# Distinctive spore architecture and developmental biology of *Turicibacter sanguinis* reveal unexpected diversity among gut spore formers

**DOI:** 10.64898/2026.01.06.697881

**Authors:** Catalina Cortés-Tapia, José García-Yunge, Ana Moya-Beltrán, Francisca Cid-Rojas, Matías Castro, C Camila Rojas-Villalobos, Fernando Gil, Raquel Quatrini, Marjorie Pizarro-Guajardo, Daniel Paredes-Sabja

**Affiliations:** ANID – Millennium Science Initiative Program – Millennium Nucleus in the Biology of the Intestinal Microbiota, Santiago, Chile; Centro Científico y Tecnológico de Excelencia Ciencia & Vida, Santiago, Chile; Departamento de Informática y Computación, Facultad de Ingeniería, Universidad Tecnológica Metropolitana, Chile; Department of Biology, Texas A&M University, College Station, Texas, USA; Programa de Doctorado en Biología Computacional, Facultad de Ingeniería, Arquitectura y Diseño, Universidad San Sebastián, Santiago, Chile; Microbiota-Host Interactions & Clostridia Research Group, Center for Biomedical Research and Innovation (CIIB), School of Medicine, Faculty of Medicine, Universidad de los Andes, Santiago, Chile; Facultad de Ciencias, Universidad San Sebastián, Santiago, Chile

**Keywords:** *Turicibacter sanguinis*, sporulation, germination, Erysipelotrichia, spore, engulfment, feeding-tube

## Abstract

Sporulation is a widespread but incompletely characterized trait among gut commensals, where it underpins microbial persistence, transmission, and ecological resilience. Most insights into spore biology derive from Bacilli and Clostridia, yet little is known about sporulation in phylogenetically distant gut-associated lineages. *Turicibacter sanguinis*, a strict anaerobe linked to host serotonin metabolism, lipid homeostasis, and neurodegenerative disease, represents one such understudied taxon. Here, we integrate ultrastructural, physiological, and comparative genomic analyses to define the sporulation and germination program of *T. sanguinis*. We show that *T. sanguinis* forms heat-resistant spores with a canonical core–cortex–coat architecture but displays previously undescribed features including a dual-layered outer envelope and bimodal electron-dense coat morphotypes. Developmental stages of sporulation follow canonical stages of *Bacillus*- and *Clostridium*-like sporulation while genomic analyses reveal a hybrid regulatory architecture combining Clostridial-type Spo0A initiation with Bacillus-like late-stage sigma factor control. Germination assays and genomic signatures further indicate a nutrient-responsive, Bacillus-like pathway involving Ger-family receptors, SpoVA-mediated Ca–DPA release, and CwlJ- and SleM-type cortex hydrolases. Together, these findings identify *T. sanguinis* as a distinct spore-forming lineage within the human gut microbiota and expand the known diversity of sporulation strategies across the Firmicutes.

## INTRODUCTION

The human gastrointestinal (GI) tract is populated with a highly diverse community of microbes (1, 2). Nearly 2000 bacterial species have been identified in the human gut, and it is estimated that the gut microbiota contains nearly 150 times more genes than the human genome (3, 4). This large genetic and metabolic potential of the gut microbiome underpins its ubiquity in nearly all aspects of human biology, including health maintenance, development, aging, and disease (5).

Throughout an individual’s lifespan, the gut microbial community is subjected to multiple episodes of dysbiosis with profound implications in its health-related roles, including the resistance to infectious diseases, metabolic balance, cognitive decline, aging, and neurodegenerative conditions such as Alzheimer’s disease (AD) further emphasizing the role of the microbiome (6-9). The most common cause of external perturbations to the microbiome are diet, medications (e.g., antibiotics), and environmental exposures (10, 11).

Recovery from these disruptions depends not only on the cessation of the perturbing factor but also on the ability of commensal microbes to recolonize the gut and re-establish functional community networks (11). Recolonization is therefore a critical determinant of both short- and long-term outcomes of microbiome disturbance. Importantly, nearly 40% of gut commensal genera are capable of forming spores (12), a feature that allows them to persist outside the host (13), withstand environmental stressors, and efficiently reseed the gut following perturbation (14). This widespread capacity for sporulation positions spore-forming commensals as key mediators of microbiome resilience and stability.

Among the spore-forming commensals, *Turicibacter sanguinis*, a Gram-positive member of the Turicibacteraceae family, represents a particularly intriguing component of the gut microbiota. This bacterium harbors a functional serotonin transporter and metabolizes gut serotonin (15), and likely modulate gut-serotonin levels with potential implications in the gut–brain axis. Notably, *Turicibacter* abundance has been reported to increase in individuals with Alzheimer’s disease (16-20), further implicating it in the interplay between dysbiosis and neurodegeneration. Moreover, extensive studies have linked *T. sanguinis* with alterations of host’s bile acid metabolism (15, 21-23). In addition to these unique physiological features, *T. sanguinis* forms resilient spores, which likely contribute to its persistence in the gut ecosystem and ability to recolonize following perturbations. Despite its potential role in gut-brain axis, little is known about *T. sanguinis* basic physiology.

In this work, we focalized on the characterization of *T. sanguinis* spore formation. Specifically, our results demonstrate that *T. sanguinis* clusters as a phylogenetically diverse group separated from members of the Erysipelotrichia class. *T. sanguinis* spore formation initiates after 2 days in YCFA agar plates, following canonical developmental stages as the well-studied *Bacillus subtilis* and *Clostridioides difficile*. Ultrastructural analysis revealed that *T. sanguinis* spores, while lacked an exosporium layer, had an outermost crust-like layer, underlined by a coat-like layer that had two distinctive types (thick and thin). Additionally, these spores contained similar spore-core DPA levels *as C. difficile* spores and germinated in the presence of culture rich medium. Genomic analysis of an assembled hybrid genome reveals that *T. sanguinis* encodes, the master regulator of sporulation, Spo0A, and all four canonical sporulation specific RNA polymerase sigma factors, with some sigma factor specific regulators (SpoVT and GerE). Notably, the germination machinery seems to include predicted GerA-family of receptors, SpoVA DPA-channel, and CwlJ-type spore peptidoglycan cortex-specific enzymes. Overall, these results provide the first insight into spores of the Erysipelotrichia class, specifically of the *Turicibacteraceae* family.

## MATERIAL AND METHODS

### Bacterial growth conditions

Bacterial microbiota strains isolated, including *T. sanguinis* K031, were routinely grown at 37 °C under anaerobic conditions (90 % N_2_, 5 % CO_2_ y 5 % H_2_) in a Bactron EZ2 anaerobic chamber (Shellab, USA) in previously described YCFA medium (12): Casitone 10 g/L (BD, USA), yeast extract 2.5 g/L (BD, USA), NaHCO_3_ 4 g/L (Merck, USA), L-cysteine 1 g/L (Merck, USA), K_2_HPO_4_ 0.45 g/L (Merck, USA), KH_2_PO_4_ 0.45 g/L (Merck, USA), NaCl 0.9 g/L (Merck, USA), MgSO_4_ x 7H_2_O 0.09 g/L (Sigma-Aldrich, Chile), CaCl_2_ 0.09 g/L (Merck, USA), resazurin 1 mg/L (Sigma-Aldrich, Chile), hemin 10 mg/L (Sigma-Aldrich, Chile), biotin 10 μg/L (Sigma-Aldrich, Chile), cobalamin (vitamin B12) 10 μg/L (Sigma-Aldrich, Chile), p-aminobenzoic acid 30 μg/L (Sigma-Aldrich, Chile), folic acid 50 μg/L (Sigma-Aldrich, Chile), pyridoxamine 150 μg/L (Sigma-Aldrich, Chile). After autoclave (121 °C, 15 min), supplemented with: sodium acetate 33 mM (Sigma-Aldrich, Chile), sodium propionate 9 mM (Sigma-Aldrich, Chile), isobutyrate 1mM (Sigma-Aldrich, Chile), isovalerate 1 mM (Sigma-Aldrich, Chile), valerate 1 mM (Sigma-Aldrich, Chile), thiamine 0.05 μg/mL (Sigma-Aldrich, Chile), riboflavin 0.05 μg/mL (Sigma-Aldrich, Chile), maltose 0.2 g/mL (Sigma-Aldrich, Chile), cellobiose 0.1 g/mL (Sigma-Aldrich, Chile) and glucose 0.2 g/mL (Sigma-Aldrich, Chile). Stock solutions were made by dissolving their components in Milli-Q water and filter sterilizing (0.2-μm pore size) prior to use.

*C. difficile* R20291 strain was grown at 37 °C under anaerobic conditions (90 % N2, 5 % CO_2_ y 5 % H_2_) on Brain Heart Infusion supplemented with 0.5% yeast extract, and 0.1% L-cysteine (BHIS) broth or 1.5% agar. *E. coli* DH5α strains were grown on Luria Bertani supplemented with 50 ug/mL chloramphenicol, 10 ug/mL tetracycline, 50 ug/mL kanamycin, or 100 ug/mL ampicillin where indicated.

Sporulation of *B. subtilis* PY79 and AZ703 was induced by the exhaustion method (24). Briefly, after 35 hours of growth in Difco Sporulation (DS) medium at 37 °C with vigorous shaking, spores were collected, washed and purified. The purification was performed using KCl 1 M, lysozyme 10 mM, NaCl 1 M, SDS 0.05% and several washes with water.

### Bacterial isolation

Fecal samples were obtained from healthy Chilean individuals within the framework of the project Millennium Nucleus in the Biology of Intestinal Microbiota. This project was approved by Comité de Bioética de la Facultad de Ciencias de la Vida, Universidad Andrés Bello, file number 013-2017. All patients enrolled in this study agreed to participate and signed an informed consent form.

Once stool samples were collected in sterile containers without preservation media (one fecal sample per donor - minimum 0.5 g), samples were reduced in an anaerobic chamber (Bactron EZ2, ShellLab) within 1 h of passing to preserve the viability of anaerobic bacteria. Culture media, sterile phosphate-buffered saline (PBS) and all other materials used for culturing were placed in the anaerobic cabinet 24 h before use to reduce oxygen levels. The fecal samples were divided into two fractions; one part was homogenized in reduced PBS (0.1 g stool/ml PBS) and was serially diluted (10-1 to 10^-8^) and plated directly onto reduced YCFA agar plates and incubated 72 - 96 h at 37 °C under anaerobic conditions. The other fraction was treated with an equal volume of 70 % (w/v) ethanol (e.g., 3 g of sample mixed with 7 mL of ethanol 100 %), for 4 h at room temperature under ambient aerobic conditions to kill vegetative cells. Cells were precipitated by centrifugation (6,500 x *g*, 10 min), washed 3 times with PBS, and the pellet was resuspended in reduced PBS under anaerobic conditions, and plated onto YCFA agar plates and incubated 24 - 48 h under anaerobic conditions at 37 °C. Their quality and morphology were evaluated by classical microbiological techniques (macro and microscopic observation). The verified colonies were propagated in YCFA broth, incubated at 37 °C for 24 h under anaerobic conditions and finally stored in DMSO (Sigma-Aldrich, Chile) 5 % at -80 °C.

### DNA extraction and 16S rRNA amplification

*T. sanguinis* isolate was identified by 16S rRNA amplification and subsequent whole-genome sequencing. Genomic DNA was phenol extracted using adapted protocols (25). Briefly, 1 mL of O/N culture of each strain isolated was centrifuged at 14,000 x *g* for 5 min, supernatant was discarded and homogenized with 12.5 μL lysozyme (10 mg/mL) (Merck) and 62.5 μL Buffer TES (Tris-HCl 50 mM (Merck), EDTA 0.5 mM (Merck), SDS 1 % (Merck) to incubate at 37 °C for 30 min with dry heat. 2.5 μL of Proteinase K (20 mg/mL, Merck) and 15 μL of SDS 10 % was added to the tube and incubated at 55 °Cfor 20 min under wet heat. Once the sample reached room temperature (22 °C), 75 μL of phenol:isopropanol:chloroform (25:24:1) were added to the sample, and hand swirled for 5 min before centrifugation at 14,000 x *g* for 5 min. The aqueous phase was transferred into a new Eppendorf tube and genomic DNA was precipitated with a solution of 250 μL of ethanol and 10 μL of NaCl (Merck) 2.5 M at -20 °C for 15 min. Finally, the sample was centrifuged at 14,000 x *g* for 10 min, supernatant was discarded, and precipitated genomic DNA was resuspended in 30 μL of ultrapure water with RNase and storage at -20 °C until use.

The 16S rRNA amplification was done with universal primers: 7F 5′-AGAGTTTGATYMTGGCTCAG-3’y 1510R 5′-ACGGYTACCTTGTTACGACTT-3′. PCR reaction was prepared with: 8 μL Mix 2X KAPA HiFi, 1 μL primer 7F, 1 μL primer 1510R, 1 μL genomic DNA and 9 μL ultrapure water to a final vol of 20 μL, and following conditions: initial denaturation of 95 °C 3 min, 35 cycles of 98 °C 30 s, 60 °C 20 s, 72 °C 2 min and a final extension of 72 °C for 2 min. PCR product was purified and sequenced by Macrogen USA. The isolate of interest was named as K031.

### Illumina and PacBio DNA sequencing

Genomic DNA was extracted from a culture of *T. sanguinis* K031 using DNeasy Blood & Tissue Kit (Qiagen) following manufacturès recommendations with the following modifications: a pellet of an overnight culture (OD_600_ of 0.3) was resuspended in 180 μL enzymatic lysis buffer and incubated for 30 min, then 25 μg/mL of RNAase were added and incubated at 37 °C for 30 min and at 56 °C for 1 h. DNA was sequenced with MicrobesNG (University of Birmingham, UK) according to the manufacturès protocol; DNA was quantified in triplicates with the Quant-iT dsDNA HS assay in an Ependorff AF2200 plate reader.

For Illumina short read sequencing, genomic DNA libraries were prepared using Nextera XT Library Prep Kit (Illumina, San Diego, USA) following the manufacturer’s protocol with the following modifications: 2 ng of DNA were used as input, and PCR elongation time was increased to 1 min from 30 s. DNA quantification and library preparation were carried out on a Hamilton Microlab STAR automated liquid handling system. Pooled libraries were quantified using the KAPA Biosystems Library Quantification Kit for Illumina on a Roche light cycler 96 qPCR machine. Libraries were sequenced on the Illumina HiSeq using a 250 bp paired end protocol.

For PacBio sequencing, Single Molecule, Real-Time sequencing (SMRT) libraries were prepared using the SMRTbell Express Template Prep Kit 2.0 (PacBio) following the manufacturer’s protocol, including DNA damage repair, end repair, and adapter ligation. Libraries were purified with AMPure PB beads and size-selected on a BluePippin (Sage Science) with a 10–15 kb cutoff to enrich long inserts. Following library QC (Qubit and TapeStation), sequencing primer was annealed and polymerase was bound (Sequel II/IIe Binding Kit), and complexes were loaded on a Sequel IIe instrument using SMRT Cell 8M. Data were collected with a 15–30 h movie time, and Circular Consensus Sequencing (CCS) reads (“HiFi”) were generated in SMRT Link using default parameters to yield Q20+ consensus reads suitable for downstream assembly and polishing.

### *Turicibacter sanguinis* K031 genome assembly and annotation

Raw reads were preprocessed using Trimmomatic v0.36 (26) to trim adaptors and to remove low quality reads. Read quality was visualized in FastQC v0.11.8 (http://www.bioinformatics.babraham.ac.uk/projects/fastqc/). *De novo* assembly of Illumina reads was performed with Velvet v1.2.10, retaining reads with quality scores > Q30. Quality control and *de novo* assembly of PacBio long reads were conducted using Canu v2.2 (27). Hybrid assembly was then performed with SPAdes v3.14.0 (28), in which PacBio long reads were used to bridge Illumina short reads. Resulting contigs were further refined using an in-house pipeline that included manual sorting, curation, and comparative genomic analyses. Assembly integrity was validated by mapping Illumina reads back to the draft genome using Bowtie2 (29). The final genome sequence was deposited in NCBI GenBank (JAGBKO000000000) and annotated using the NCBI Prokaryotic Genome Annotation Pipeline (PGAP) (30) (**Table S1A**). The *dnaA* gene was designated as the first locus of the final assembly.

### Publicly available *Turicibacter spp*. Database

Additional publicly available genomes of *Turicibacter* spp. were retrieved in February 2024 using NCBI Datasets command-line tools (https://www.ncbi.nlm.nih.gov/datasets/) with the parameters: taxon “*Turicibacter*” and --assembly-source “GenBank” (**Table S1B**). Retrieved genomes were assessed for completeness (> 90%) and duplication (< 5%) using BUSCO and universal orthologue analysis as described by Raes et al. (31). Assembly statistics for the curated dataset of 31 genomes were summarized (**Table S1C**).

### Taxonomic allocation using 16S rRNA and ribosomal protein phylogenies

A total of nine 16S rRNA genes identified in the hybrid *T. sanguinis* K031 genome sequence were compared and clustered according to sequence identity (**Table S2A**). The taxonomic allocation of *T. sanguinis* K031 isolate was verified based on 16S rRNA gene sequence analysis. Ribosomal RNA genes were downloaded from the NCBI nucleotide database using Entrez Direct v14.6 (https://www.ncbi.nlm.nih.gov/books/NBK179288/; February 2024) with the query: “*Turicibacter* 16S ribosomal RNA”. A total of 359 sequences were extracted and filtered for >1000 bp in length. An additional 320 sequences (>1000 bp) were extracted from GenBank genome assemblies of the *Turicibacter* genus. This yielded a dataset of 431 sequences, which were clustered at 100% identity using CD-HIT v4.8.1 (32), with minimum alignment coverage of 100%. A total of 59 representative sequences were selected for downstream taxonomic and phylogenetic analyses (**Table S2B**). The 16S rRNA gene of isolate K031 (Table S2A) was aligned against this curated panel using BLASTN (NCBI BLAST+; megablast; E-value ≤ 1×10⁻²⁰, identity ≥ 97%, query coverage ≥ 98%) (32, 33), and assignments were confirmed by a reciprocal BLAST in which each top subject sequence was queried back against the K031 dataset; annotations were accepted only when the reciprocal best hit returned K031 with comparable identity and coverage. Because of continuous taxonomic reclassification of the Erysipelotrichales order, to select appropriate outgroups, we followed the NCBI taxonomy to the order level, and from there chose two families, each represented by one genus and one type strain genome with complete sequence, ensuring that none were candidate taxa.

Sequences were aligned with MAFFT v7.490 using the FFT-NS-2 algorithm, and the resulting alignment was trimmed with trimAl v1.2 at a gap threshold of 0.5. Phylogenetic relationships were inferred using Maximum Likelihood (ML), Bayesian Inference (BI), and Neighbor-Joining (NJ). The ML tree was constructed with PhyML v3.0 (34). The BI tree was generated using MrBayes v3.2.7, run for 10,000 generations with trees sampled every 100 generations, and posterior probabilities were calculated after discarding the first 25% of sampled trees. The NJ tree was constructed with QuickTree (35) using the UPGMA and Kimura options, and bootstrap support values were calculated from 10,000 iterations. An additional phylogenetic tree was generated with FastTree v2.1.10 (double precision) (36) and visualized with the Interactive Tree Of Life (iTOL) v4 (http://itol.embl.de). Well-supported nodes were defined as those with bootstrap values ≥95%.

In parallel, a concatenated set of ribosomal proteins (**Table S2C**) was aligned with MAFFT v7.310 using the L-INS-i method (37). Alignments were checked and refined manually. Phylogenetic trees were again reconstructed using three approaches. ML trees were generated with PhyML v3.0 (34) using Smart Model Selection (SMS v1.8.4) and 1000 bootstrap replicates. NJ trees were built with FastME v2.0 (38) using the WAG substitution model with a gamma distribution and 1000 bootstrap replicates. Finally, BI trees were generated with MrBayes v3.2.6 (39), run for 1000 generations with trees sampled every 100 generations, and posterior probabilities were calculated after discarding the first 30% of trees as burn-in, using a nucmodel and MCMC (Markov chain Monte Carlo) analysis.

### Overall genome relatedness indexes calculations

The specific assignment of K031 (and related strains) was further validated using whole-genome relatedness indexes, namely Average Nucleotide Identity (ANI) (40, 41) and digital DNA–DNA Hybridization (dDDH) (42, 43) using the 31 *Turicibacter* spp. genomes (**Table S1C**). The overall genome relatedness indexes (OGRIs) were calculated from all possible pairwise genome comparisons using both nucleotide- and amino acid–based metrics. Average nucleotide identity based on BLAST (ANIb) was calculated using the Python package pyani (https://github.com/widdowquinn/pyani) (41) (**Table S3A**); *in silico* DNA-DNA hybridization index (dDDH) was calculated using the Genome-to-Genome Distance Calculator with recommended formula 2 (43) and species cutoff limits defined by Meier-Kolthoff and colleagues (42) available at http://ggdc.dsmz.de (**Table S3B**). Additionally, average amino acid identity (AAI) was calculated using CompareM (https://github.com/donovan-h-parks/CompareM) (**Table S3C**).

### Growth profile of *T. sanguinis* K031

The growth profile of *T. sanguinis* K031 was determined by inoculating 1% (v/v) of an overnight culture into fresh YCFA broth. Cultures were incubated anaerobically at 37 °C for 12 h. At 0, 2, 4, 5, 6, 7, 8, 9, 10, 11, and 12 h, 1 mL aliquots were collected and optical density at 600 nm (OD_600_) was measured. Non-inoculated medium served as a blank control, and these baseline values were subtracted from sample measurements. The assay was performed three independent times, each in triplicate, and results are presented as mean ± SEM, with growth curves generated using Prism 8 software (GraphPad).

### Spore preparation and purification

*Turicibacter sanguinis* K031 spores were purified by adapting previously described *Clostridioides difficile* spore purification protocols (44, 45). Briefly, 100 μL of a 1:100 dilution of an overnight culture in YCFA broth was spread on YCFA agar plates (1.5% [w/v] Bacto agar; BD, USA) and incubated anaerobically at 37 °C for 3 days. Colonies were scraped from plates into ice-cold sterile Milli-Q water and washed five times by centrifugation at 14,000 × g for 5 min at 4 °C. The pellet was resuspended in 45% (w/v) autoclaved Nycodenz (Axell, USA) and centrifuged at 14,000 × g for 40 min to separate spores. The spore-containing pellet was recovered and washed five additional times with ice-cold sterile Milli-Q water (14,000 × g, 5 min) to remove residual Nycodenz. Lastly, spores were enumerated using a Neubauer chamber, adjusted to a final concentration of 5 × 10^9^ spores/mL, and stored at −80 °C.

### Transmission electron microscopy (TEM)

*T. sanguinis* spores were processed and analyzed using prior published methods (44, 45). Briefly, three sample were prepared: (i) 1 mL of overnight culture of *T. sanguinis* K031 grown in YCFA broth, (ii) 1 mL of sporulated culture grown on YCFA plates, and (iii) 100 μL of purified spores from frozen stock (5 × 10^9^ spores/mL). Samples were centrifuged at 14,000 × g for 5 min, the supernatant was removed, and pellets were fixed by gently layering 500 μL of 3% glutaraldehyde in 0.1 M cacodylate buffer (pH 7.2) without resuspension. Pellets were incubated at 4 °C for at least 24 h (primary fixation), followed by postfixation in 1% osmium tetroxide in 0.1 M cacodylate buffer (pH 7.2). Samples were then washed in cacodylate buffer, stained with 1% tannic acid for 30 min, and dehydrated through an acetone gradient (30% for 30 min, 50% for 30 min, 70% overnight, 90% for 30 min, and twice in 100% for 20 min each). Dehydrated pellets were infiltrated with Spurr’s resin using graded acetone:resin ratios of 3:1, 1:1, and 1:3 (40 min each), resuspended in 100% Spurr’s resin for 4 h, and polymerized overnight at 65 °C. Ultrathin sections (90 nm) were cut with a microtome, mounted on glow-discharged carbon-coated grids, and double-stained with 2% uranyl acetate and lead citrate. Sections were imaged using a Philips Tecnai 12 BioTWIN transmission electron microscope at the Electron Microscopy Facility of the Pontificia Universidad Católica de Chile.

### India ink staining

Five microliters of *T. sanguinis*, *C. difficile*, and *B. subtilis* spores from frozen stock (5 × 10^9^ spores/mL) were diluted 1:100 in sterile Milli-Q water and mixed with an equal volume of India ink (Sigma-Aldrich, Chile), as previously described (47). Samples were mounted on 1.7% agarose pads (46) and examined by phase-contrast microscopy as described by Shuster et al. (2019) for visualization of the characteristic spore halo.

### Fluorescence microscopy of sporulation stages

An aliquot of 100 μL from a 1:100 dilution of an overnight culture of *T. sanguinis* K031 in YCFA broth was spread onto YCFA agar plates and incubated anaerobically at 37 °C for 12, 24, and 48 h. After incubation, 2 mL of sterile Milli-Q water was added to each plate, and colonies were scraped and transferred to microcentrifuge tubes under anaerobic conditions. Samples were centrifuged at 3,780 × g for 10 min, the supernatant was removed, and the volume was adjusted to 1 mL with sterile Milli-Q water. For fluorescence staining, 50 μL of culture collected at 12, 24, or 48 h was centrifuged at 7,000 × g for 5 min, and the pellet was resuspended in 50 μL of staining solution containing 10 μg/mL FM 4-64 (phospholipid stain, Thermo Fisher, Chile), 9 μM MitoTracker Green (MTG, protein stain, Thermo Fisher, Chile) and 4.5 μM Hoechst (DNA stain, Thermo Fisher, Chile). Samples were incubated at room temperature for 2 min in the dark, centrifuged at 3,780 × g for 5 min, and the supernatant was discarded. Pellets were resuspended in 20 μL of 1X PBS, mounted on 1.7% agarose pads (46), and imaged by fluorescence microscopy using a QImaging camera with an exposure time of 100 ms. Fluorescently stained morphotypes observed during sporulation were visualized and merged using ImageJ, images were organized according to canonical sporulation stages described in spore-forming bacteria (10, 12), and morphotypes of each stage were quantified and plotted using Prism 8 software (GraphPad).

### Dipicolinic acid (DPA) assay

DPA content of spore cores was quantified as previously described (48). Briefly, 5 μL of purified *T. sanguinis* spores (5 × 10^9^ spores/mL) were diluted 1:200 in 1X TBS buffer and denatured by heating at 100 °C for 30 min. Samples were centrifuged at 14,000 × g for 5 min, and supernatants were collected (49). Aliquots of 125 μL were transferred to a 96-well plate and supplemented with 800 μM TbCl₃. The Ca-DPA concentration was determined by monitoring the fluorescence of the DPA–Tb³⁺ complex (excitation 270 nm, emission 545 nm) using a Synergy H1 Hybrid Multi-Mode Reader (BioTek) (48). Assay was performed in biological triplicate, each with technical triplicates.

### Spore colony formation efficiency

The colony formation efficiency of *T. sanguinis* spores was evaluated using freshly purified spores (0 days of storage) and spores stored at 4 °C for 2, 4, 7, or 28 days. Aliquots of spores (1 × 10^7^ spores/mL) were plated on YCFA agar with or without prior heat activation (50, 60, or 70 °C for 20 min) and incubated anaerobically at 37 °C for 24 h. Spore viability was calculated as [(c.f.u./mL)/ (spore particles/mL)] x 100 and expressed relative to wild-type strain.

### Spore germination

An aliquot of 5 µL of a stock of 5x10^9^ of purified *T. sanguinis* spores per mL was heat-activated at 60 °C for 20 min. Non-activated spores were used as control. Both, heat- and non-heat-activated spores were incubated in 1 mL of 70:30 and BHIS, culture media typically used for *Clostridia* and *Bacillus* (50) spore germination, and YCFA, previously verified as germination media for *T. sanguinis* spores. OD_595_ of each sample was measured at 30-, 60-, 120- and 240-min using 1X PBS as optical density control. Additionally, after 4 hours of incubation, aliquots were analyzed in an agarose pad under the phase contrast microscope, and phase-bright, -gray, and - dark spores counted (n = 200 spores were counted per treatment condition).

### Ortholog identification pipeline for sporulation and germination related proteins

The predicted proteome of *T. sanguinis* K031 (JAGBKO000000000) was functionally annotated against KEGG/KO, COG, Pfam, and NCBI Conserved Domain Database (CDD). Ortholog identification proceeded in 2 stages. The first stage was to compile a curated dataset of hallmark sporulation, germination and spore-structure proteins from five spore-former model species *B. subtilis* strain 168 (NC_000964), *B. cereus* ATCC14579 (NZ_CP034551), *B. anthracis* Ames ancestor (NC_007530), *C. difficile* strain 630 (NC_008262) and *C. perfringens* SM101 (NC_009089) (**Table S4**). Each reference protein was mapped to functional profiles (KEGG Orthology, COG, Pfam, and NCBI Conserved Domain Database) (**Table S5**). Homology relationships were evaluated using BLASTP and HMMER (hmmsearch). Proteins were retained as candidates when they matched the same KO/COG/Pfam/CD signature as a reference or produced a significant HMM hit to the corresponding profile (default thresholds: E-value ≤ 1e-5 for BLASTP and hmmsearch; alignment coverage ≥ 50% of the reference/domain; domain architecture consistent with the reference model). All candidates were manually curated using gene-neighborhood and operon context and synteny with model organisms; final calls are reported in Supplementary Table S6.

## RESULTS

### Gene and genomic evidence confirm K031 as a *Turicibacter sanguinis* representative

During the construction of our gut microbiota collection, serially diluted fecal samples from healthy donors were plated into YCFA plates, and individual colonies subjected to full-length 16S rRNA gene amplicon sequencing with universal primers 7F–1510R, and identified one of the isolates as *Turicibacter sanguinis*. This isolate was whole genome sequenced to construct a high-quality hybrid genome of *T. sanguinis* K031 was obtained (3.3 Mb; 34.4 % G+C) and assembled into eight scaffolds (N50 = 2,145,658) at > 99x depth of coverage from 129.9 Mb of Illumina and 20.3 Gb of PacBio raw reads (**Table S1A**). The graft genome sequence was deposited at the GeneBank database under the accession IDs JAGBKO000000000.

Nine 16S rRNA gene copies were identified in the *T. sanguinis* K031 genome.Clustering of these copies resolved two allelic 16S rRNA gene groups: eight identical sequences (Group 1) and a single allele with minor divergence (J3478_02830; > 98% identity to Group 1) (**Table S2A**). Phylogenetic analyses incorporating representatives of both of these groups, together with reference publicly available *Turicibacter* 16S rRNA sequences and carefully curated outgroups (**Table S2B**), consistently placed strain K031 within the *Turicibacter sanguinis* clade (**Figure S1A**). These results confirm the assignment of K031 to *T. sanguinis* species.

To generate a more comprehensive phylogeny of *T. sanguinis* and enable direct comparison of strain K031 with other sequenced strains and meta-assembled genomes tentatively assigned to the species, we retrieved a set of phylogenetically informative ribosomal proteins (RPs) (**Table S2C**) (52). Concatenated alignments of seven universally present RPs (RP7) were analyzed using Bayesian inference (BI), Maximum Likelihood (ML), and Neighbor-Joining (NJ) methods. Whereas ML and NJ yielded trees with relatively unstable support (**Figure S1B, S1C**), BI produced a robust and well-supported phylogeny (**Figure 1A**). In all three trees, strain K031 consistently clustered within the *T. sanguinis* clade together with the type strain MOL361^T^ (53) (**Figure 1A, S1B, S1C**). Unclassified *Turicibacter* strains, including one recently reassigned to *T. bilis* (54), formed a broader sister clade (**Figure 1A**).

**Figure 1.**
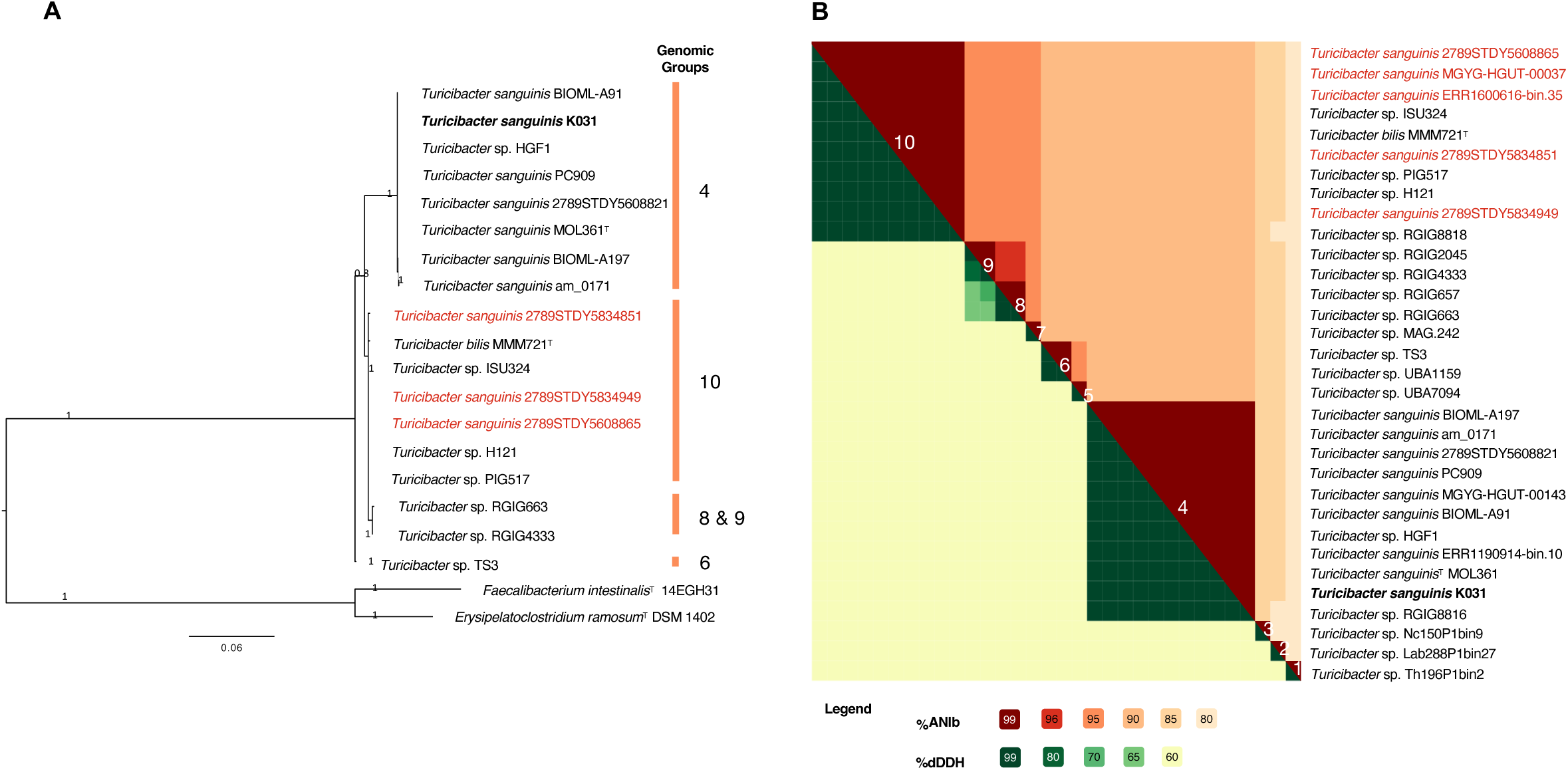
Phylogenetic and genome relatedness analysis of *Turicibacter sanguinis K031*. **(A)** MCC Bayesian Phylogenetic tree with a concatenated alignment of 7 ribosomal proteins present in all *Turicibacter* sp. strains and outgroups in the analysis (<u>Table S2C</u>). (**B**) Heatmap of Average Nucleotide Identity or ANIb (%, orange scale) and the DNA-DNA Hybridization index or dDDH (%, green scale). Genomic groups in both panels are numbered similarly (from 1 to 10). Thresholds used for species delimitation are the following: ANI > 96% (same genomic species); dDDH > 70% (same genomic species). Type strains are marked with a T superscript. *Faecalibacterium intestinalis*^T^ 14EGH31 and *Erysipelatoclostridium ramosum*^T^ DSM 1402 were used as outgroups.

ANI and dDDH whole-genome relatedness analysis between K031 strain and publicly available *Turicibacter* spp. genome assemblies (as of February 2024, Table S1), yielded concordant results, resolving 10 genomic groups above species-level relatedness thresholds (**Figure 1B**). These analyses confirmed the genomic coherence of *T. bilis* (Group 10) and *T. sanguinis* (Group 4), with strain K031 unambiguously placed within *T. sanguinis* (Group 4). Beyond these named taxa, several additional groups appear, which are represented by a single genome and lack support from the ribosomal-protein phylogeny (**Figure 1**), indicating that their taxonomic status remains provisional pending additional sampling.

### *T. sanguinis* cellular morphology suggests S-layer

To begin characterizing the basic physiology of *T. sanguinis*, we investigated its growth kinetics. An overnight culture was diluted 1:100 into pre-warmed YCFA broth and incubated anaerobically at 37 °C. The exponential growth phase lasted for the first 4 h, reaching a maximum OD_600_ of 0.44, which remained stable during the subsequent 8 h of monitoring (**Figure 2A**). After 16 h, however, cultures appeared to undergo lysis, consistent with the weak Hoechst staining observed (**Figure S2A**). Planktonic cells from 8 h cultures were examined by phase-contrast microscopy (**Figure 2B**) and transmission electron microscopy (**Figure 2C–E**). Phase-contrast imaging confirmed a bacillary morphology (**Figure 2B**), while TEM revealed that cell division proceeded by symmetric binary fission (**Figure 2C**). TEM also showed numerous spherical and rod-shaped profiles (**Figure 2C, 2D**), which likely reflect cross-sectional orientations of cells in thin resin sections. At higher resolution, TEM further suggests that the cell wall of *T. sanguinis* has an outer S-layer–like structure surrounding the peptidoglycan layer and underlying cytoplasmic membrane (**Figure 2E**).

**Figure 2.**
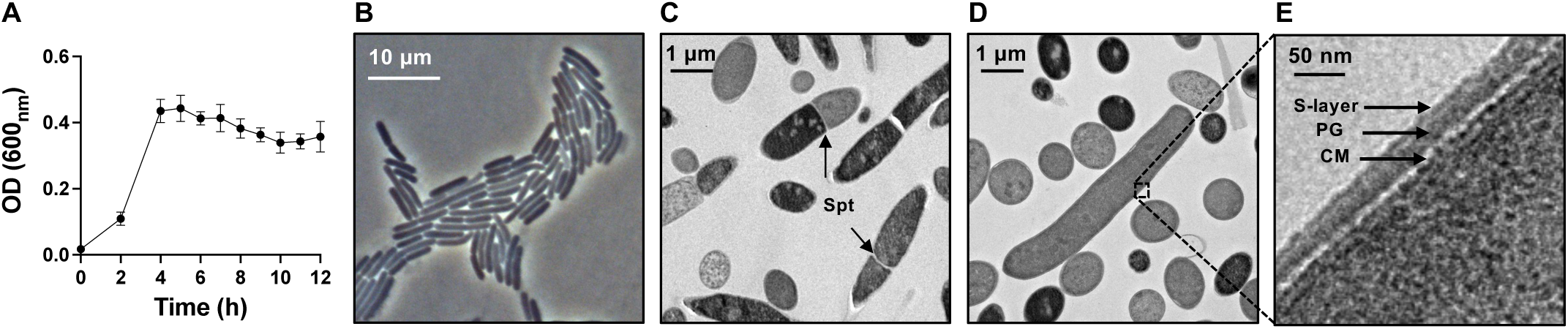
Vegetative growth of *T. sanguinis* K031 in YCFA broth. (**A**) Growth curve of *T. sanguinis* K031 after 2, 4, 6, 8, 10 and 12 h of incubation in YCFA broth at 37 °C under anaerobic conditions. Error bars indicate the standard error of the mean (S.E.M.). The experiment was done three times with technical triplicates. (**B**) Phase contrast microscopy imaging of an 8 h YCFA culture of *T. sanguinis* K031. (**C**), (**D**) and (**E**) represent transmission electron micrograph images of an 8 h YCFA culture of *T. sanguinis* K031 with a magnification of 3,400x. (**C**) Arrows denote the symmetric septum (spt) formation during cell division. Panels D and E, represent vegetative cells, where (**E**) is the zoom of bacilli’s surface from (**D**). (**D**) Spherical morphologies represent a transversal cut section of *T. sanguinis* bacilli. (**E**) S-layer, Peptidoglycan (PG) and cell membrane (CM) are highlighted with black arrows. Bar scales are highlighted.

### *T. sanguinis* forms phase-bright spores

Among the different colony morphologies, *T. sanguinis* formed flat colonies with a white-irregular-shiny viscous texture, and a wavy margin, with a bubble on the center of the colony (**Figure 3A**). A notable observation when culturing *T. sanguinis* on YCFA agar plates was the appearance of phase-bright structures resembling dormant spores under phase-contrast microscopy (**Figure 3B, S2B**), which contrasted with absence of sporulating cells in YCFA broth cultures (**Figure 2B, S2A**). To investigate the sporulation dynamics of *T. sanguinis*, overnight YCFA broth cultures were diluted 1:100, plated onto YCFA agar, and incubated anaerobically for 12, 24, and 48 h at 37 °C. Phase-contrast microscopy revealed the appearance of sporulating cells (25 %) and phase-bright free spores (∼13 %) as early as 12 h of incubation, and after 48 h phase bright free spores reached ∼85 % of total (**Figure 3C**).

**Figure 3.**
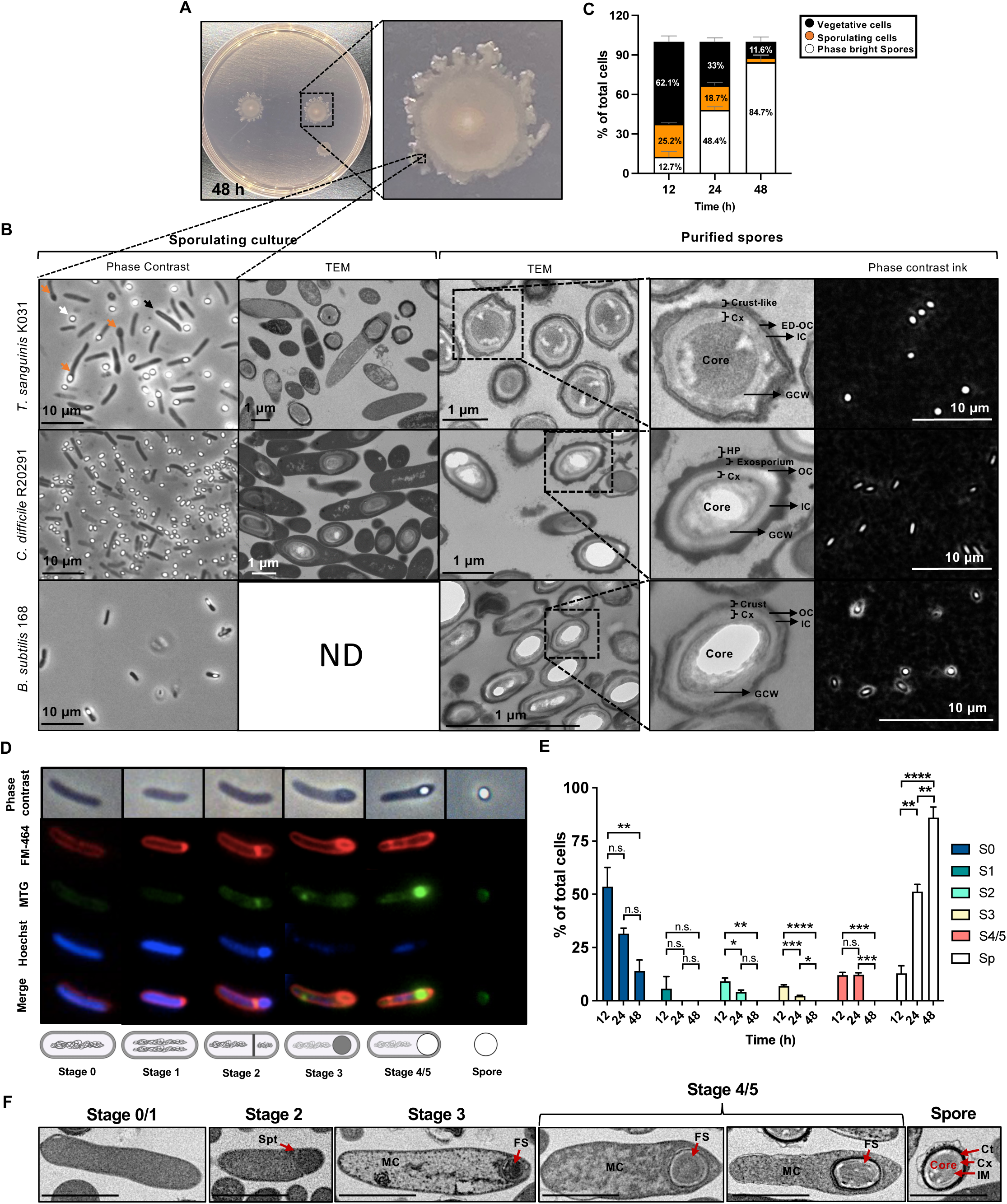
Ultrastructural and physiological characterization of *T. sanguinis* K031 sporulation. (**A**) Representative *T. sanguinis* K031 48 h colony in YCFA agar plate. (**B**) Visualization of sporulating cultures and purified spores of *T. sanguinis*, *C. difficile* and *B. subtilis* by phase contrast microscopy and transmission electron microscopy. First two left panels represent phase contrast microscopy and TEM images of *T. sanguinis* K031, *C. difficile* R20291 and *B. subtilis* strain 168 sporulated cultures after 48 h of anaerobic incubation at 37°C on YCFA, 70:30 or LB plates for each strain, respectively. Black, orange, and white arrows denote vegetative cell, sporulating cell and free spore respectively. Subsequent two panels correspond to TEM images from purified spores, and individual zoomed spore of *T. sanguinis* K031, *C. difficile* R20291 and *B. subtilis* strain 168 harvested. In the zoom of individual spores are pointed out each structural layer. Core, germ cells wall (GCW), cortex (Cx), inner coat (IC), outer coat (OC) or electron-dense outer coat (ED-OC), exosporium, hair-like projections (HP) and crust or crust-like. Panels further to the right evaluates presence of crust in purified spores of *T. sanguinis*, *C. difficile* R20291 and *B. subtilis* using India ink (presence of polysaccharides). *B. subtilis* strain 168 and *C. difficile* strain R20291 spores were positive and negative control for halo formation around the spore respectively. TEM images of sporulating culture of *T. sanguinis* K031 and *C. difficile* R20291 were taken at a magnification of 3,400x and 11,500x respectively. TEM images of purified spores of *T. sanguinis* K031, *C. difficile* R20291 and *B. subtilis* strain 168 were taken at a magnification of 6,700x, 20,500x and 60,000x respectively. Bar scales are highlighted. (**C**) Phase contrast microscopic quantification of sporulation efficiency of *T. sanguinis* K031 during 12, 24 and 48 h of incubation at 37 °C under anaerobic conditions. Average from three independent experiments and error bars indicate the standard error of the mean (S.E.M.). (**D**) Sporulation developmental stages in *T. sanguinis*. Representative micrographic images of 12, 24, and 48h of YCFA agar plate sporulating cultures of *T. sanguinis* were stained with Hoechst (DNA), FM4-64 (membrane-impermeable dye) and MTG (membrane-permeable dye). Schematic development and examined by phase contrast and fluorescence microscopy. Schematic representation of the developmental stages is depicted below selected micrographs. (**E**) Frequency of developmental stages at various time points in YCFA agar plates. Error bars indicate the standard error of the mean (S.E.M.). The graph represents three independent experiments. Statistical analysis was performed using One-way ANOVA followed by Tukey’s multiple comparison test, ns: no significance, p < 0.5: *p < 0.1, **p < 0.01, ***p < 0.001, ****p < 0.0001. (**F**) Transmission of electronic micrographs at a magnification of 3,400x of 48 h sporulation culture of *T. sanguinis* K031 in YCFA plates. Representative images of the different developmental stages are shown MC: mother cell, FS: forespore, Spt: septum, Ct: coat, Cx: cortex, IM: inner membrane. Scale bars represent 2 µm.

To further characterize sporulation, we compared *T. sanguinis* sporulating cultures with those of *Clostridioides difficile* strain R20291 and *Bacillus subtilis* 168 using phase-contrast and TEM (**Figure 3B**). Phase contrast microscopy and electron micrographs demonstrate that *T. sanguinis* forms endospores similarly as *C. difficile* and *B. subtilis* (**Figure 3B**). TEM analysis showed that *T. sanguinis* spores possess the same general structural organization as those of *B. subtilis* and *C. difficile*: a spore core surrounded by an inner membrane, a germ cell wall (GCW), a thick cortex peptidoglycan layer (Cx), inner (IC) and outer (OC) coat layers, and an outermost layer resembling the *B. subtilis* crust rather than the exosporium observed in *C. difficile* spores (**Figure 3B**).

In *B. subtilis* and its close relatives (referred as the *B. subtilis* group), the outermost crust or exosporium layer is adorned with polysaccharides (PS), where sugars like rhamnose are common in these species (55). A traditional method to identify the presence of polysaccharides in the spore surface is using India ink, which is unable to penetrate through a PS layer, thus producing a white halo around the spores (47). Therefore, since electron micrographs suggest that the outermost structure of the *T. sanguinis* spores is similar to the crust observed in *B. subtilis*, we assayed *T. sanguinis* spores with India ink to test whether the *T. sanguinis* sporés outermost layer would also contain polysaccharides as those of the *B. subtilis* crust. However, as we can observe in Figure 3C, unlike the halo observed in *B. subtilis* spores, neither *T. sanguinis* nor *C. difficile* spores exhibited a halo surrounding the spore (**Figure 3B**). These results suggest that although the morphology of the crust of *T. sanguinis* spores is similar to that of *B. subtilis,* it might not be adorned with polysaccharides. Despite this slight biochemical difference, given the ultrastructural similarities of the outermost layer of *T. sanguinis* and *B. subtilis* spores, the outermost layer of *T. sanguinis* spore will be named as “Crust-like” outermost layer.

### Developmental stages of sporulation in *T. sanguinis* follow canonical models

Given that *T. sanguinis* forms similar spores to those from model spore-formers *B. subtilis* and *C. difficile,* as was mentioned before (**Figure 3C**), we next asked whether its sporulation process also followed the canonical sequence described in Bacilli and Clostridia. To address this, *T. sanguinis* was grown under sporulating conditions on YCFA plates, the medium in which spore formation had previously been observed. Because phase-bright spores were already evident after 12 h of incubation, we monitored sporulation at 12, 24, and 48 h under anaerobic conditions at 37 °C using transmission electron microscopy and fluorescence microscopy with Hoechst, MTG, and FM4-64 staining to visualize DNA, proteins, and membranes, respectively (**Figure 3D**). Based on these analyses, we identified five sporulation stages. Stage 0 (S0) represented vegetative cells, while Stage 1 (S1) showed increased Hoechst staining consistent with chromosome duplication. Stage 2 (S2) was characterized by asymmetric septum formation at one pole of the cell, stained with FM4-64 and MTG, indicating its lipid and protein composition. Stage 3 (S3) corresponded to an engulfed forespore, evidenced by FM4-64 staining of the surrounding membrane Stage 4/5 (S4/5) were presented by a phase-grey and phase-bright endospore within the mother cell, respectively, with intense MTG (**Figure 3D, 3F**). To quantify the distribution of morphotypes over time, we scored cells at each stage after 12, 24, and 48 h of incubation (**Figure 3E**). While the relative abundance of S0 decreased after 48 h of sporulation, stage S1 remained constant, whereas stages S2, S3, S4/5 decreased similarly as S0 after 48 h of sporulation (**Figure 3E**). In the case of free spores, these increased up to ∼80 % of the population after 48 h (**Figure 3E**). These results demonstrate, that *T. sanguinis* sporulation is asynchronous and progresses from vegetative growth toward spore formation over time in YCFA plates, following the canonical stages of sporulation (S1–S5).

### Architecture of *T. sanguinis* spores

TEM analysis of sporulating *T. sanguinis* cultures revealed that endospores within mother cell sporangia displayed either a thick or thin electron-dense (ED) surface layer (**Figure 4A**), suggesting the presence of two distinct spore morphotypes, as previously described in *C. difficile* (44). Interestingly, upon measuring purified *T. sanguinis* spores by transmission electron micrographs, we observed that the entire spore population could be divided into thin or thick outer surface spores (**Figure 4B, S3**). Quantification of two independently prepared batches of *T. sanguinis* spores demonstrated that depending on the spore preparation, 35 to 83 % of the spores had a thick surface layer spores (**Figure 4C**), denoting potential phase variations in this trait.

**Figure 4.**
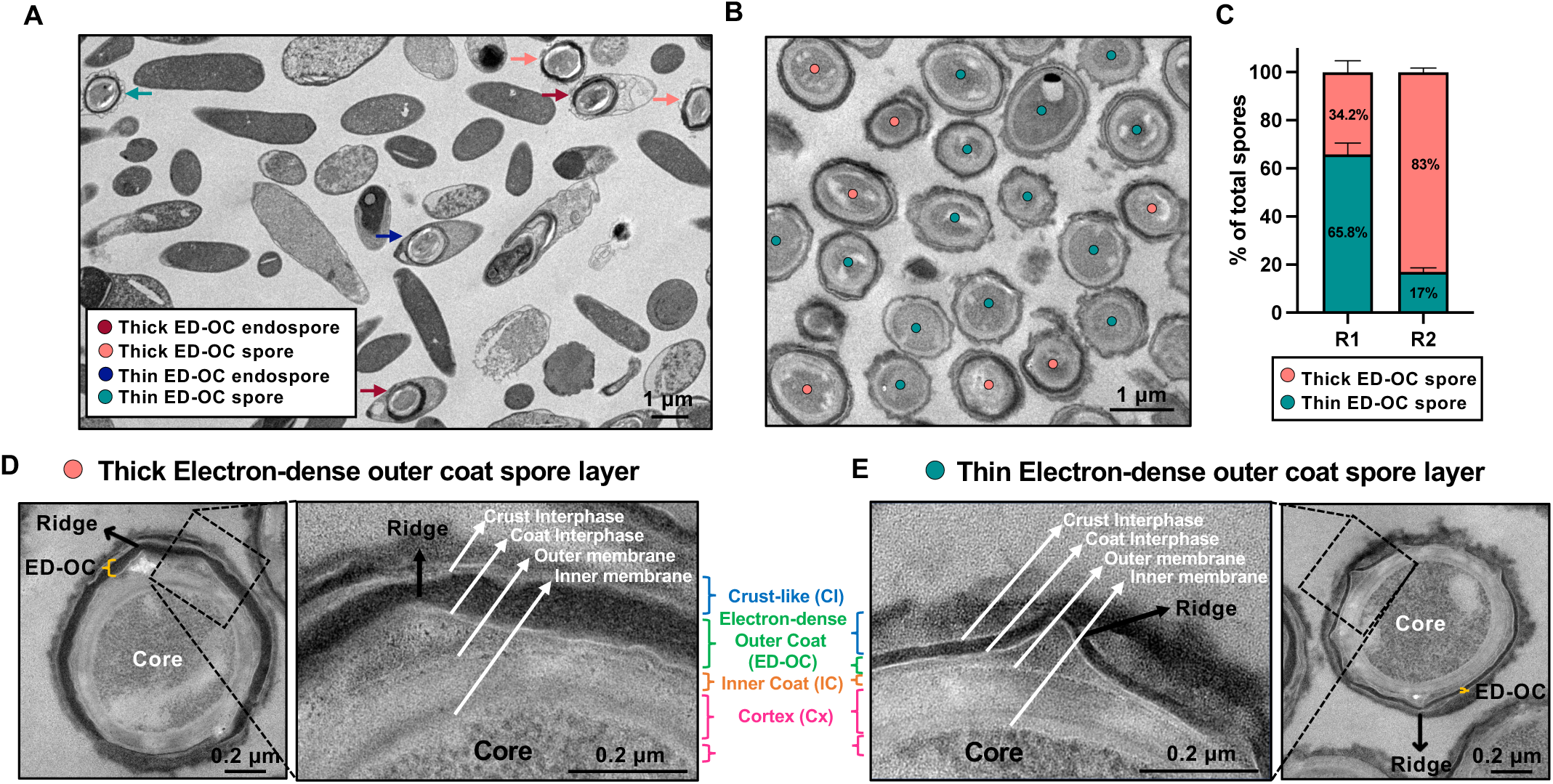
Transmission electron microscopy reveals thick and thin *T. sanguinis* K031 spore morphotypes. (**A**) Electron micrographs of *T. sanguinis* K031 sporulated culture, growth for 48 h in YCFA plate, with a magnification of 3,400x. Thin and Thick ED-OC endospores and spores are highlighted. (**B**) Representative TEM image of *T. sanguinis* K031 purified spores with a magnification of 5,300x. *T. sanguinis* spores with a thick (coral circles) or thin (green circles) electron-dense outer coat (ED-OC) layer are highlighted. (**C**) Quantification of thick and thin ED-OC spores respect to the total spore population. A total of 335 spores were visualized from 2 biological replicates of purified spores (R1: n = 221; R2: n = 114). Error bars indicate the standard error of the mean (S.E.M.). Bars represent percentage of thick (coral) and thin (green) ED-OC spores in population. (**D** and **E**) Zoom of thick (Panel D) and thin (Panel E) ED-OC spores. The distinctive spore layers, membranes and interphases are shown. Ridge, Crust-like (Cl), Crust interphase, Electron-dense outer coat (ED-OC), Coat interphase, Inner coat (IC), Outer membrane, Cortex (Cx), Germ cell wall (GCW), Inner membrane and Core.

At higher magnification (36,000×), thick-type spores exhibited a prominent ED layer (**Figure 4D**), whereas this layer was much thinner in thin-type spores (**Figure 4E**). Despite this difference, both morphotypes shared a similar internal organization: a spore core bounded by an inner membrane, surrounded by a cortex-like peptidoglycan layer (Cx), an outer membrane, and multiple concentric coat layers including the inner coat (IC), the ED outer coat (ED–OC), and an intermediate coat layer (**Figure 4D, 4E**). Notably, the first layer external to the outer membrane lacked the lamellae-like features characteristic of *B. subtilis* and *C. difficile* spores (13, 44, 56, 57).

A striking feature in both morphotypes was the presence of surface ridges originating from the layer beneath the ED–OC (**Figure 4D, 4E**). Ridge counts followed an average of nine ridges per spore. Surrounding the ED–OC was a diffuse outermost layer resembling the *B. subtilis* crust, separated by a defined crust interphase in both type-spores (**Figure 4D, 4E**). Altogether, these observations indicate that *T. sanguinis* spores exhibit a hybrid architecture, sharing structural features of both, *B. subtilis* and *C. difficile* spores, and, as in *C. difficile*, forming two distinct morphotypes characterized by thin or thick electron-dense outer layers.

### Analysis of *T. sanguinis* spore architecture reveals thick/thin spores and an outer crust

Ultrastructural observations revealed that *T. sanguinis* produces two distinct spore morphotypes characterized by either a thin or thick electron-dense outer coat (ED-OC). To determine whether these differences followed a structured pattern within the population, we performed a quantitative morphometric analysis of spore ultrastructure (**Figure 5A-C**). For whole spore diameter and spore core (SC) size measurements, each spore was divided into three symmetrical axes through the center, and diameters were averaged (**Figure 5A**). Measurements were performed on both thin and thick ED–OC spores, which were color-coded for comparative analysis (green, thin; coral, thick) (**Figure 5D**). To assess the thickness of structural layers beyond the core, including the cortex (Cx), inner coat (IC), ED–OC, and crust-like (CL) layers measurements were made along lines drawn through ridges and inter-ridge regions, averaging six per spore (**Figure 5C**). Ridge height was calculated as the difference between maximum and minimum IC thickness.

**Figure 5.**
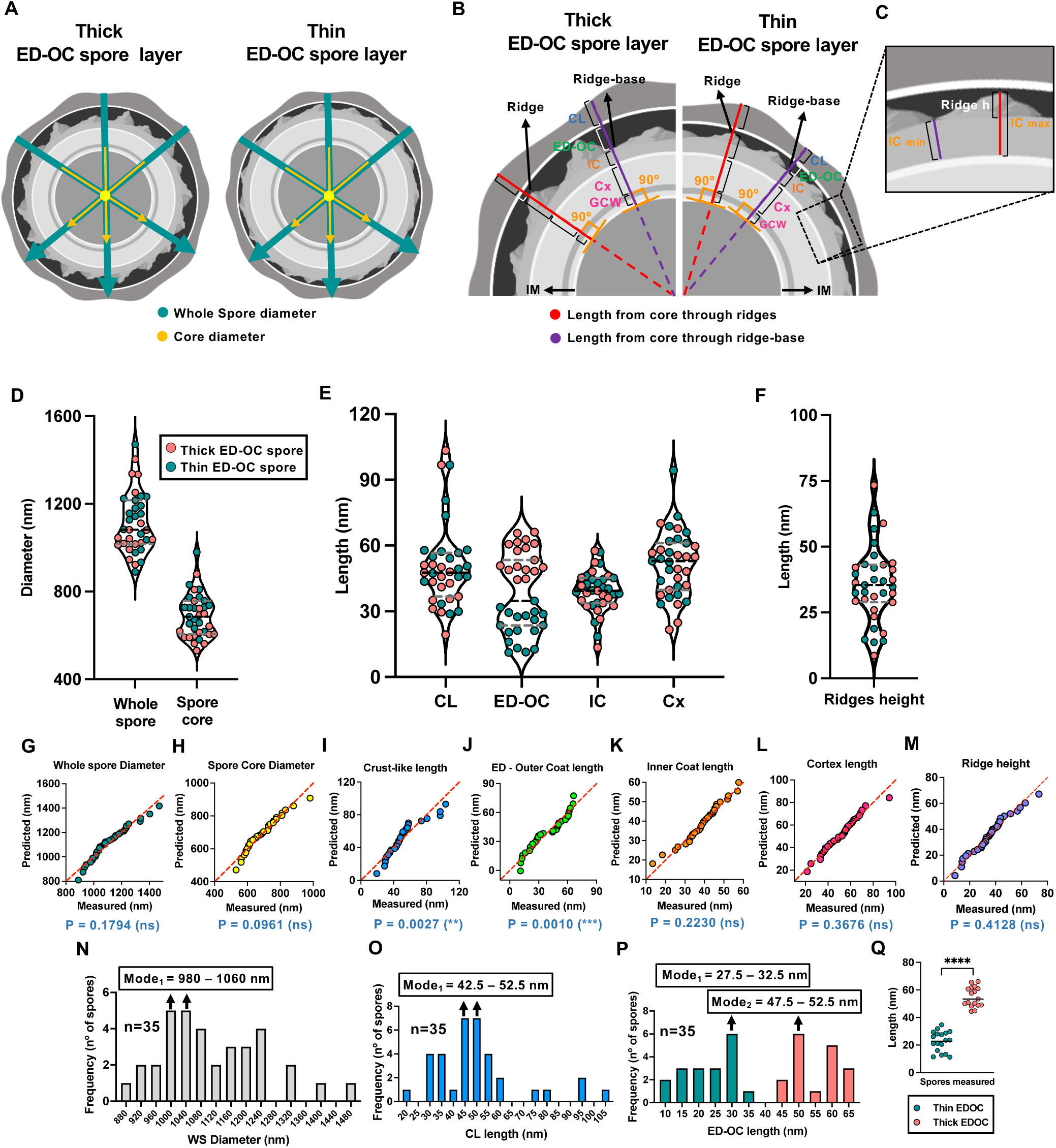
*T. sanguinis* K031 spore quantitative analysis of ultrastructural layers. (**A**) Schematic representation of measurement of whole spore length quantification in both ED-OC morphotypes in *T. sanguinis*. First, 3 concentric vectors are drawn in each spore to measure Whole spore (green arrows) and Spore Core (yellow arrows) diameters. Measurements for each spore and spore core diameter represents the average of three measurement per spore. (**B**) Model representation of thin and thick ED-OC spores’ structural layers length’s measurement. First, six lines are drawn to measure spores Cortex (Cx), Inner coat (IC), Outer coat (ED-OC) and Crust-like (Cl) layers. 3 lines crossing from the top of the ridge (red line), and 3 lines crossing the bottom of the ridges (purple line). Measurements for each spore represents the average of six measurement per spore. (**C**) Model representation of ridge height, where height for each ridge was measured by as the difference between the lengths between the maximum (red line) and minimum (purple line) length of the Inner coat (IC). (**D-F**) Layer-length quantification of a total of 35 individual *T. sanguinis* spores from 2 biological replicates (R1: n = 22; R2: n = 13). The length of whole spore and spore core (Panel D), Crust-like layers (CL), Outer coat (ED-OC), Inner coat (IC), and Cortex (Cx) (Panel E), and of the ridge height (Panel F) was quantified. In D-F, thick and thin spores are represented by coral and green circles, respectively. Black and gray dashed lines represent median, and quartiles of spores measured respectively. (**G-M**) QQ plots of the normal distribution of the length of the diameters of Whole spore (**G**) and Spore core (**H**) diameters, the length of Crust-like layer (**I**), Electron dense outer coat (**J**), Inner coat (**K**), Cortex (**L**) and high of Ridge (**M**). Plots were tested for normal distribution by D’Agostino & Pearson test; p < 0.05 is indicative of departure from normality. Each circle symbol represents the average length of one spore. (**N-P**) Histograms of frequency distribution of the length of whole spore (**N**), Crust-like (**O**) and ED - Outer coat (**P**) were plotted using Prism 8 software and Modes for skewed (**O**) and bimodal distributions (**P**) were highlighted. Green and coral bars from panel P represent Thin and Thick ED-OC spores respectively. (**Q**) Graph representing thin (green circles) and thick (coral circles) ED-OC *T. sanguinis* spores’ populations. Statistical analysis was performed by two-tailed paired Student’s t test, **** p < 0.0001.

Quantitative analysis showed that *T. sanguinis* spores had an average whole spore diameter of ∼1.1 μm (range: 0.9 – 1.5 μm) and an SC diameter of ∼0.7 μm (0.5–0.9 μm) (**Figure 5D**). The CL averaged 51 nm (19–103 nm), the ED–OC 38 nm (11–66 nm), the IC 39 nm (13–58 nm), and the cortex 51 nm (22–94 nm) (**Figure 5E**). Ridge height averaged 36 nm (9–73 nm) (**Figure 5F**). These values are comparable to those reported for *C. perfringens* and *C. difficile* spores (44, 58). Inspection of thickness distributions revealed high heterogeneity across layers. Normality testing (QQ plots) showed that whole spore diameter, spore core diameter, IC thickness, cortex thickness, and ridge height followed normal distributions (**Figures 5G, 5H, 5K–M**). In contrast, CL and ED–OC thicknesses deviated from normality (**Figures 5I, 5J**). The WS histogram confirmed a clear normal distribution (**Figure 5N**), whereas the CL histogram was skewed, with a mode between 42.5–52.5 nm (**Figure 5O**), which was not observed in the other layers (**Figure S4A-D**). Strikingly, ED–OC thickness exhibited a bimodal distribution with two distinct modes (27.5–32.5 nm and 47.5–52.5 nm), suggesting two subpopulations of spores (**Figure 5P**). When spores were classified by morphology, thin and thick ED–OC spores each followed normal distributions (**Figure S4E, S4F**). Comparative analysis confirmed that thin ED–OC spores had a significantly reduced thickness (mean = 22.8 nm) compared to thick ED–OC spores (mean = 54.9 nm; P < 0.0001) (**Figure 5Q**). Importantly, ED–OC thickness did not correlate with whole spore diameter or other layer measurements (**Figure S5**), suggesting that assembly of the ED– OC is governed by a distinct regulatory mechanism, independent of overall spore size.

### *T. sanguinis* spores germinate in presence of nutrients

DPA is a hallmark of bacterial spore core that contributes to DNA protection through dehydration (57). Therefore, to assess whether *T. sanguinis* spores also contained DPA, purified spores were heat-treated and released DPA quantified fluorometrically via TbCl₃-complexation assay. *T. sanguinis* spores contained ∼11 µg/mL DPA, comparable to the ∼14 µg/mL measured in *C. difficile* R20291 spores used as a positive control (**Figure 6A**), indicating that *T. sanguinis* spores shares a canonical marker of bacterial spores.

**Figure 6.**
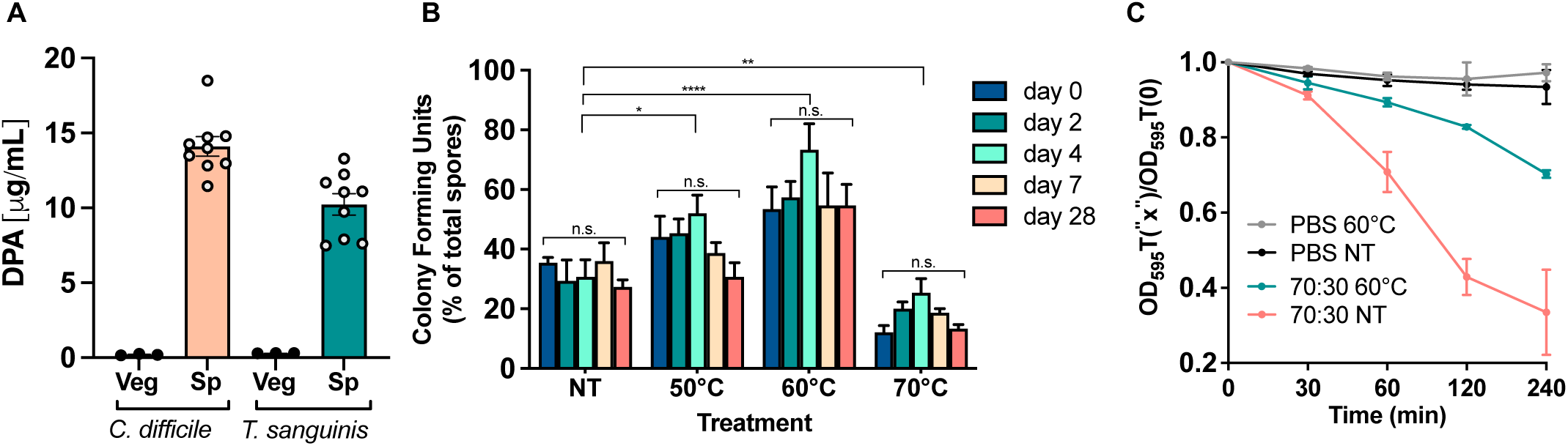
DPA content and spore germination of *T. sanguinis* spores. (**A**) Dipicolinic acid (DPA) content was quantified from vegetative cells and spores of *T. sanguinis* and *C. difficile*. (B) Colony forming units expressed was quantified by plating untreated or heat activated (50, 60 or 70 °C) spores (10^7^ spores / mL) onto YCFA agar plates and expressing values relative to total quantified spores. (**C**) Spore Germination of *T. sanguinis* K031 on 70:30. *T. sanguinis* spores were heat activated at 60°C and compared against non-activated spores (NT). Spore germination was measured by optical density. All experiments were done three independent times in three technical replicates. Error bars represent the standard error of the mean. *p < 0.05; **p < 0.01; ****p < 0.0001.

Heat activation of bacterial spores synchronizes them to germinate homogeneously and increases the spore colony forming efficiency (59). However, spores also age and loss viability though extended storage (45). Therefore, we examined how spore age and heat activation affected spore colony viability. For this, spores stored at various time points up to 28 days at 4 °C were treated with several heat activation temperatures. We observed that from all heat activation conditions, 30 min at 60 °C led to the highest colony forming efficiency of up to ∼70 % (**Figure 6B**). By contrast, spore age had no effect in pore colony forming efficiency (**Figure 6B**). Notably, untreated spores had a spore colony forming efficiency of ∼35%, indicating that while heat activation might be useful *in vitro*, under physiologically relevant conditions, heat activation of *T. sanguinis* spores is not required.

The ability of spores to return to a vegetative cells is essential for their biological function and require specific germinant(s) to trigger the process (59). We assessed whether *T. sanguinis* would germinate in commonly used nutrient rich broth such as 70:30 (typically used to sporulate *C. difficile*) or BHI (for growth of *C. difficile*) (60, 61). Hence, heat-activated and untreated spores were incubated in YCFA, 70:30, and BHIS broth, and OD_600_ was measured over 4 h (**Figure 6C and S6A**). Unexpectedly, non-heat-treated spores showed greater OD decline than heat-treated spores, particularly in YCFA medium, which exhibited an OD_600_-decrease of ∼ 40% reduction (**Figure S6A**). Phase-contrast microscopy revealed three phenotypes (phase-bright, grey, and dark spores), with quantification showing medium-dependent differences (**Figure S6B, S6C**). In YCFA, ∼50% of spores reached the grey stage but only ∼30% became fully dark. In contrast, 70:30 medium yielded ∼75% dark-phase spores within 4 h. BHIS supported minimal germination, with only 10% grey spores and 2% dark spores, regardless of heat treatment (**Figure S6C**). Altogether, these results indicate that the kinetics of germination of *T. sanguinis* spores does not require heat activation, and that nutrient rich broths are sufficient to trigger germination of *T. sanguinis* spores.

### Sporulation pathway in *T. sanguinis*

To gain insight into the sporulation pathway, we utilized a pipeline for sporulation-related orthologues discovery in *T. sanguinis* (**Figure S7**). Comparative genomic analysis revealed the absence of orthologues of the *Bacillus*-type phosphorelay components, the phosphotransfer proteins (e.g., Spo0F and Spo0B) and orphan histidine kinases (e.g., KinA-E) (**Figure 7A, Table S6**). However, key regulators of sporulation onset were identified, including *sigH* (J3478_09480) and *spo0A* (J3478_15840), along with three *sigA* paralogs (J3478_13700, J3478_14895, J3478_09755), the first of which corresponds to the *B. subtilis sigA* ortholog (**Figure 7C, 7D, S8**; **Table S6**). A *spo0E* (J3478_13625) and *ptpA*-like phosphatase (J3478_05720) orthologues, known to inhibit *spo0A* activity in *C. difficile* (62-64), were also identified (**Figure 7C, 7D; Table S6**), suggesting that *T. sanguinis* is also subjected to similar sporulation repression as observed in these species. Altogether, these genomic features indicate that while *T. sanguinis* conserves the central regulatory logic of Spo0A-mediated sporulation, the molecular activation may be unique to this lineage.

**Figure 7.**
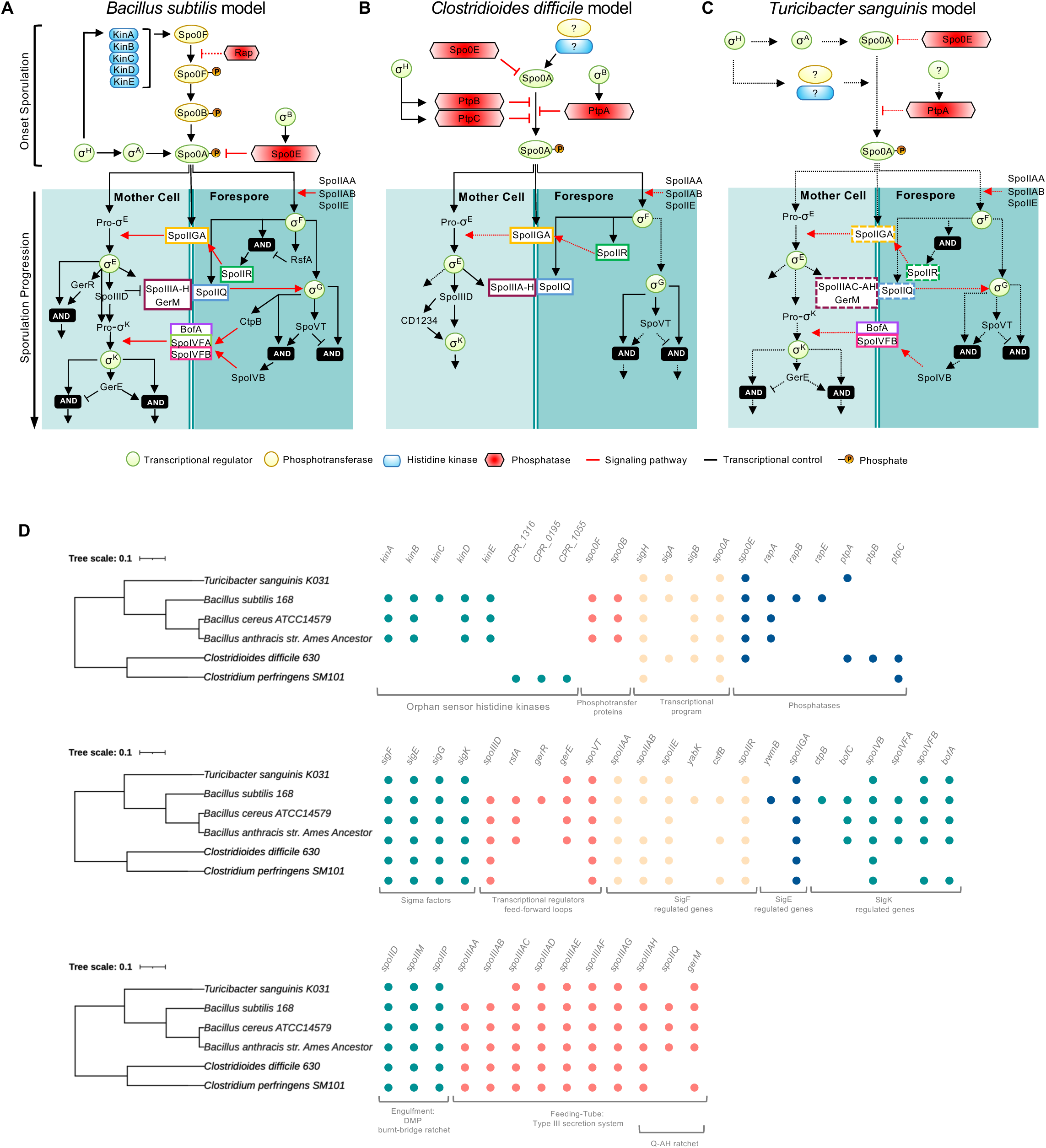
Putative sporulation pathway in *T. sanguinis*. Schematic transcriptional regulation of sporulation in *B. subtilis* (**A**) and *C. difficile* (**B**) and suggested model of *T. sanguinis* (**C**), adapted from Shen *et al*. 2019 (68). Temporal progression of sporulation in the three species are shown from top to bottom. Transcriptional regulators, phosphotransferases, histidine kinases and phosphatases from onset sporulation are enclosed in green, yellow, light blue and red boxes, respectively. As described in symbology, black arrows represent transcriptional control of gene expression, red arrows indicate signaling pathways and dashed lines indicate that the regulatory relationship remains unknown. Proteins enclosed in white boxes with colored outline directly participate in signaling between the mother cell and forespore. (**D**) Phylogenetic tree was made with 16S ribosomal RNA gene sequence (16S rRNA) from *T. sanguinis* K031, Bacilli (*B. subtilis* strain 168, *B. cereus* ATCC14579, *B. anthracis* Ames Ancestor), and Clostridia (*C. difficile* strain 630 and *C. perfringens* SM101), using ClustalW (https://www.genome.jp/tools-bin/clustalw) for their alignment. The tree was constructed using FastTree v2.1.8 with default parameters and then graphically visualized with the web tool Interactive Tree Of Life V3 (http://itol.embl.de). *B. subtilis* strain 168, *B. cereus* ATCC14579, *B. anthracis* Ames Ancestor, *C. difficile* strain 630 and *C. perfringens* SM101 genomes were downloaded from NCBI genome database with accession number NC_000964, NZ_CP034551, NC_007530, NC_008262 and NC_009089 respectively. Phylogenetic profile shows the conservation of gene presence for relevant functional categories.

Once Spo0A is phosphorylated, sporulation proceeds through activation of the four conserved sporulation-specific sigma factors, σ^F^, σ^E^, σ^G^, and σ^K^ (**Figure 7A**), which drive compartment-specific transcription in the forespore and mother cell (65-67). Genomic analysis identified orthologs of all four sigma factors in *T. sanguinis*, σ^F^ (J3478_01570), σ^E^ (J3478_00215), σ^G^ (J3478_00210) and σ^K^ (J3478_06915) (**Figure 7C, 7D, S9-S11; Table S6**). Consistent with σ^F^ regulation, the anti-sigma factors SpoIIAA (J3478_01580) and SpoIIAB (J3478_01575) were also detected (**Figure 7C, 7D, S12; Table S6**). The presence of these regulatory elements aligns with the sporulation stages observed by fluorescence and electron microscopy (**Figure 3D, 3E**) and indicates that *T. sanguinis* encodes the canonical sigma-factor cascade associated with early sporulation.

The activation of sporulation-specific sigma factors requires cross-compartment signaling between the forespore and mother cell (**Figure 7A**) (68). Genomic analysis of *T. sanguinis* revealed orthologs of several proteins involved in these signaling pathways. Components of the forespore-to–mother cell signaling axis were represented by *spoIIGA* (J3478_00220), which in Bacilli and Clostridia model organisms, is activated by σ^F^-dependent SpoIIR to trigger σ^E^ maturation (69-71). Likewise, orthologs of the σ^E^-dependent mother cell–to–forespore signaling system, including *spoIIQ–AH* (J3478_07145 and J3478_04710) and *gerM* (J3478_13875) (72-75), were identified (**Figure 7C, 7D, S13; Table S6**). In contrast, genes encoding the mother cell regulators controlling σ^K^ activation in *B. subtilis* but absent in *C. difficile*, while SpoIVFA was not detected, BofA (J3478_05245) and SpoIVFB (J3478_02455) were present in *T. sanguinis* genome (**Figure 7A, 7B, S14B, S14C; Table S6**). These findings support the presence of conserved machinery associated with activation of the late forespore transcriptional program, including σ^G^.

After asymmetric division, sporulation proceeds with engulfment of the forespore by the mother cell (68). Genomic analysis of *T. sanguinis* K031 revealed homologs of the canonical engulfment machinery (76-78), *spoIIP* (J3478_07105), *spoIID* (J3478_04715), and *spoIIM* (J3478_15865) (**Figure 7D, S15; Table S6**), indicating conservation of the septal peptidoglycan degradation system involved in this morphogenetic step.

The transcriptional program governing sporulation is modulated by σ-factor–associated feed-forward loops (79). In *T. sanguinis*, orthologs of the late-stage regulators SpoVT (σ^G^-dependent; J3478_08120) and GerE (σ^K^-dependent; J3478_13865) were identified (**Figure 7C, 7D, S16; Table S6**), whereas orthologs of early feed-forward regulators associated with σ^F^ and σ^E^ were not detected. These findings suggest that *T. sanguinis* retains regulatory fine-tuning during late forespore and mother-cell stages of development.

### Germination model of *T. sanguinis* spores

Genomic analysis of *T. sanguinis* K031 identified two tricistronic germinant receptor operons, *gerAABC* (J3478_03110–03120) and *gerKBAC* (J3478_03230–03220) (**Figure 8C**, 8D**, S17; Table S6**), consistent with the presence of nutrient-responsive germinant receptors (13, 57, 59, 80, 81). The genome also encoded four SpoVA proteins (25, 48, 49, 82-85), SpoVAC, SpoVAD, SpoVAE, and SpoVAF (J3478_01565, J3478_01560, J3478_01555, J3478_01550), organized as a tetracistronic operon (**Figure 8C**, 8D**, S18B; Table S6**), indicating conservation of the SpoVA channel responsible for Ca-DPA release during germination.

**Figure 8.**
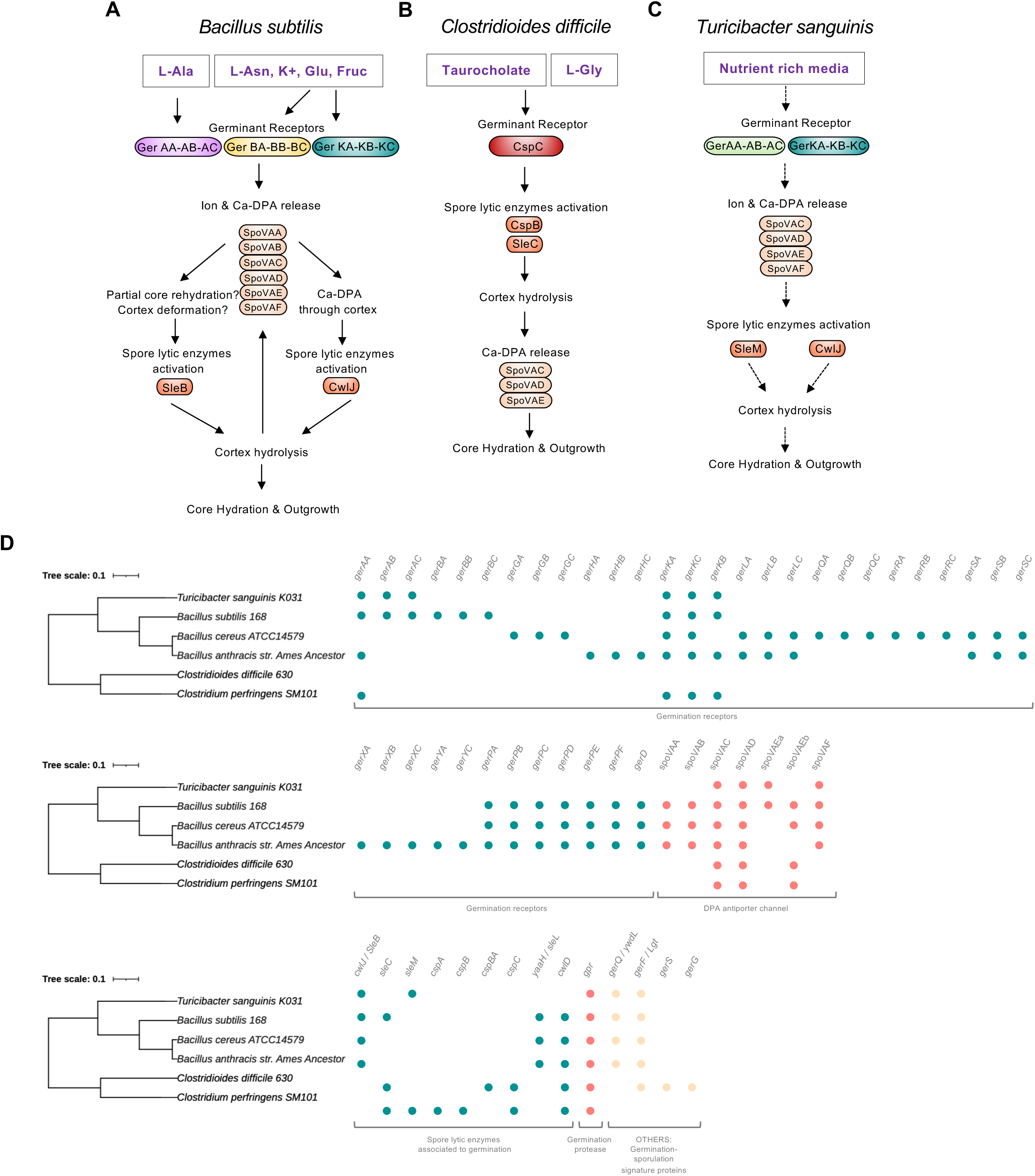
Putative germination model of *T. sanguinis* spores. Schematic spore germination signaling pathway in *B. subtilis* (**A**) and *C. difficile* (**B**) and suggested model of *T. sanguinis* (**C**), adapted from Shen *et al*. 2019. The germinating molecules that have been reported in *T. sanguinis* K031, *B. subtilis* and *C. difficile* are found in white boxes, followed by their receptor, GerA, GerB, GerK for *B. subtilis* and CspC for *C. difficile*. The suggested germination receptor of *T. sanguinis* are GerA and GerK. Proteins involved in Ca-DPA release (SpoVA) and spore lytic enzymes are shown in beige and orange cylinders. The suggested signaling pathway of *T. sanguinis* shows more similarity with *B. subtilis* model respect to sharing germination receptors GerA and GerK, and the spore lytic enzyme CwlJ for cortex hydrolysis. SpoVA is present in the three models. The order of hydrolysis of the cortex and DPA release via SpoVA is different in both models and it is unknown what could be the order during the germination of spores of *T. sanguinis*. Figure was modified/adapted from Shen *et al*. 2019 (68). (**B**) Phylogenetic tree was made with 16S ribosomal RNA gene sequence (16S rRNA) from *T. sanguinis* K031, *B. subtilis* strain 168, *B. cereus* ATCC14579, B. anthracis Ames Ancestor, *C. difficile* strain 630 and *C. perfringens* SM101, using ClustalW (https://www.genome.jp/tools-bin/clustalw) for their alignment. The tree was constructed using FastTree v2.1.8 with default parameters and then graphically visualized with the web tool Interactive Tree Of Life V3 (http://itol.embl.de). *B. subtilis* strain 168, *B. cereus* ATCC14579, *B. anthracis* Ames Ancestor, *C. difficile* strain 630 and *C. perfringens* SM101 genomes were downloaded from NCBI genome database with accession number NC_000964, NZ_CP034551, NC_007530, NC_008262 and NC_009089 respectively. Phylogenetic profile shows the conservation of gene presence for relevant functional categories.

Although no homologs of *C. difficile* Csp proteins or the SleC cortex hydrolase were detected, *T. sanguinis* encoded orthologs of the *C. perfringens* SleM hydrolase (J3478_03615) and the *B. subtilis* cortex lytic enzyme CwlJ (J3478_08195) and GerQ (J3478_08200) involved in CwlJ cortex-localization (**Figure 8C**, 8D**, S18A, S19B, S19C; Table S6**). Two additional spore germination proteins were also found, GerF (J3478_10960) and Gpr (J3478_07110) (**Figure S19A, S19D**). Together, these findings indicate that *T. sanguinis* possesses a germination genetic composition like *B. subtilis* and other nutrient-germinating Bacilli than to the bile-salt– responsive germination system of *C. difficile*.

## DISCUSSION

The gut microbiota has profound implications for human health, and perturbations in its composition can transiently or permanently deplete beneficial commensals (11). Among the diverse gut ecosystem, *T. sanguinis*, a member of the *Turicibacteraceae* family, represents an intriguing commensal organism. Its abundance has been reported to increase in Alzheimeŕs disease (16-20) and has been linked to alterations in host bile acid metabolism (15, 21-23). Notably, *T. sanguinis* has also been suggested to modulate intestinal serotonin levels, with potential implications for the gut-brain axis communications (15). Nearly 40% of gut microbiota genera have the potential to form spores, highlighting sporulation as a key mechanism for persistence, both within and outside of the host environment (12, 13). Most bacterial spores studied to date belong to the class of Bacilli and Clostridia. In this context, our study expands understanding of *T. sanguinis* sporulation and spore physiology, providing detailed insight into its encoded sporulation and germination machinery. Furthermore, we enrich the repertoire of spore ultrastructural studies within the *Turicibacteraceae* family by presenting the first comprehensive ultrastructural analysis of *T. sanguinis* spores.

A major contribution of this work is the demonstration that *T. sanguinis* forms heat-resistant spores similarly as the phylogenetically distant lineages of model spore formers. Unlike the well-studied spore-former *B. subtilis*, *T. sanguinis* exhibited asynchronous sporulation, resembling the pattern observed in *C. difficile* (13, 57, 86). Yet, in contrast to *C. difficile* (87), *T. sanguinis* sporulation was highly efficient, reaching > 80% sporulation on solid medium after 48 h. The resulting spores displayed a characteristic spherical morphology with a diameter of ∼1.1 μm, distinct from the more elongated spores of Bacilli and Clostridia species (58, 88, 89). Although *T. sanguinis* spores show only mild to moderate heat resistance compared with species *C. difficile* or *C. perfringens* with highly resistant spores (58), their ability to withstand elevated temperatures with limited loss of viability indicates adaptation to survival outside the host. This level of heat tolerance, together with efficient sporulation on solid substrates, supports the idea that *T. sanguinis* can persist in environmental reservoirs and be transmitted between hosts via spore-mediated routes, adding an important ecological dimension to its role as a gut commensal and potentially contributing to the spread of health (10). Overall, these observations warrant further investigation into the role of *T. sanguinis* spores in its life cycle and host colonization dynamics.

Transmission electron microscopy revealed that the architecture of *T. sanguinis* spores shares similar canonical features observed in *C. difficile* and *B. subtilis* (56, 57), but with distinct differences in the outermost layers. In *T. sanguinis*, we identified a dehydrated core densely packed with ribosomes, surrounded by an inner membrane, a germ cell wall, and a spore-specific PG cortex resembling those of well-characterized endospores (56). However, *T. sanguinis* spores displayed striking differences in the outer layers compared with *C. difficile* and *B. subtilis*. The inner coat of *B. subtilis* spores is approximately 75 nm thick, lamellar, and lightly stained in TEM micrographs (91), whereas *T. sanguinis* exhibited a thinner (13–58 nm) inner coat lacking lamellar organization but presenting prominent ridge-like structural features. Although similar ridge-like patterns have been reported in *Bacillus subtilis* (92), the more organized and uniform ridges observed in *T. sanguinis* may reflect lineage-specific architectural adaptations, possibly reflecting multiple initiation, termination sites of coat assembly, or specialized regions of structural reinforcement.

A striking observation in *T. sanguinis* spores is the presence of two distinct outer-layer morphotypes, thin and thick ED–OC spores, each following a normal distribution and independent of overall spore size. In addition, the inner coat comprises two distinguishable sublayers, with the innermost layer displaying regularly spaced ridge-like structures across the entire surface, a unique architecture not observed in *B. subtilis* or *C. difficile* spores (57, 91). To our knowledge, this represents the first report of simultaneous formation of thick and thin coat-layer phenotypes within a single bacterial population during sporulation. Moreover, the bimodality of the ED-OC layer is similar to reports in *C. difficile*, where spores exhibit thin or thick exosporium layers (13, 44, 45, 86). As in *C. difficile*, ED–OC thickness in *T. sanguinis* was independent of whole-spore diameter, reinforcing the idea that its formation is governed by dedicated regulatory mechanisms distinct from those controlling global spore morphogenesis. Notably, the drastic change in proportion of thick spores in both replicates, suggest that regulation of this phenotype is variable in nature. In *C. difficile*, bimodal exosporium layers have been associated with differences in tropism toward host molecules such as E-cadherin, fibronectin, and vitronectin (93, 94), contributing to functional diversification within the spore population. Similarly, the presence of both thin and thick ED–OC morphotypes in *T. sanguinis* might reflects a regulated developmental program rather than stochastic variation in coat assembly that contributes to functionally specialized spore populations in *T. sanguinis*. Future comparative proteomic and genetic studies will be necessary to determine whether the thin and thick ED–OC morphotypes of *T. sanguinis* correspond to functionally specialized subpopulations, as observed in *C. difficile* (13, 44, 45, 86).

Another contribution is that the outermost layer of *T. sanguinis* spores displayed an amorphous appearance reminiscent of the crust observed in *B. subtilis* spores (47), that surrounds the underlying ED-OC layer. Its molecular composition appears to differ from that of the *B. subtilis* crust, as *T. sanguinis* spores did not exhibit a halo when stained with India ink (**Figure 3**), a hallmark of the polysaccharide-rich crust in *B. subtilis* (47). To date, two types of outermost structures have been described surrounding the spore coat: (i) the amorphous “crust” layer in *B. subtilis* (47, 92), and (ii) the exosporium, characterized by its hair-like projections and, in some cases, a distinct interspace separating it from the underlying coat, as observed in *B. anthracis*, *B. megaterium*, and *C. difficile* (57). The organization of the crust as a double-layered in *T. sanguinis* expands the known diversity of spore envelopes. Ongoing work in our laboratory aims to identify the molecular components and regulatory pathways governing the assembly of these outer layers.

The genomic features underlying sporulation in *T. sanguinis* reveal that this organism employs a regulatory logic that blends elements of both *Bacillus*- and *Clostridium*-like developmental systems. Although *T. sanguinis* initiates sporulation efficiently, its genome lacks the hallmark *Bacillus*-type phosphorelay components, including Spo0F, Spo0B, and the KinA–E kinases, indicating that the classical relay is not required to activate Spo0A. Instead, the presence of Spo0A-inhibitory phosphatases (Spo0E and a PtpA orthologue) suggests a regulatory strategy more closely aligned with Clostridia, where Spo0A phosphorylation is controlled directly rather than through multistep phosphotransfer (68, 95-98). This hybrid architecture highlights an evolutionary divergence in how early sporulation decisions are regulated within the Erysipelotrichia.

Despite differences in initiation, downstream elements of the sporulation program appear highly conserved. The presence of all four sporulation sigma factors (σ^F^, σ^E^, σ^G^, and σ^K^), along with their corresponding regulatory proteins, suggests that the compartmentalization of transcription between the forespore and mother cell in in *T. sanguinis* follows the classical Bacilli framework (68, 99-101). Genes associated with engulfment and intercompartment communication—such as *spoIIP*, *spoIID*, *spoIIM*, *spoIIIQ*, and *germ*, further support conservation of mid-developmental morphogenesis (66, 68). Notably, however, only the late feed-forward loops (σ^G^/SpoVT and σ^K^ /GerE) appear to be retained, whereas earlier regulatory modules typical of Bacilli (σ^F^/RsfA, σ^E^/SpoIIID, σ^E^/GerR) were not detected. This reduction of early network complexity may reflect lineage-specific streamlining of transcriptional control.

Regulation of σ^K^ provides another key point of divergence. While *C. difficile* employs a simplified system in which σ^K^ is active upon translation (66, 79), *T. sanguinis* encodes BofA, SpoIVB, and the SpoIVFB protease, suggesting that σ^K^ activation depends on proteolytic processing of pro-σ^K^, as seen in Bacilli (68). Although early events resemble Clostridia models, later steps in coat formation and maturation appear to rely on Bacilli-like mechanisms. Together, these features indicate that *T. sanguinis* integrates regulatory strategies from both classical endospore formers, providing a unique evolutionary perspective on sporulation within the Erysipelotrichia and raising important questions about how these mixed systems are coordinated during development.

The genomic and phenotypic characteristics of *T. sanguinis* suggest that its germination mechanism is more closely aligned with Bacilli-like pathways than with those of Clostridia. The presence of two complete tricistronic Ger receptor operons (*gerKABC* and *gerAABC*) implies that nutrient-derived signals are key triggers for germination (59, 68, 102). Given that Ger-family receptors in *B. subtilis* recognize amino acids and sugars such as L-asparagine, D-glucose, and L-alanine (AGFK pathway) (80), *T. sanguinis* may respond to analogous small molecules, although this remains to be experimentally validated. A second hallmark of Bacilli-like germination is evident in the organization of SpoVA proteins associated with Ca–DPA transport, which is consistent with the presence of Ca-DPA in *T. sanguinis* spores. The arrangement of all SpoVA components in a single operon suggests coordinated control of Ca–DPA release, a process that typically precedes cortex hydrolysis in Bacilli (80). The predicted reliance on CwlJ- and SleM-like hydrolases further supports this interpretation, as these enzymes fulfill the cortex-lytic role played by SleB/SleC in *Bacillus* or by the CspB–SleC system in *C. difficile* (57, 68). Taken together, the genomic evidence strongly points toward a germination mechanism in which nutrient sensing triggers Ca–DPA efflux through SpoVA, followed by cortex degradation— mirroring the temporal hierarchy described for *B. subtilis* (103).

Functional assays reinforce this interpretation. The detection of ∼10 µg/mL of DPA places *T. sanguinis* at the lower end of the DPA spectrum reported for well-characterized spores (104, 105), confirming that DPA sequestration is a conserved feature of its developmental program. Furthermore, the strong enhancement of germination after heat activation parallels responses described for *C. difficile* and *B. subtilis* (106-108), indicating that heat-induced conformational priming is conserved across these phylogenetically distant spore-forming lineages. The differential germination response to YCFA and 70:30 media suggests that the germinant component(s), are present in both media, but absent or at suboptimal concentrations in BHIS, serve as germinants for *T. sanguinis*, an idea that warrants further biochemical investigation. Altogether, these findings position *T. sanguinis* among the Bacilli-like germinating taxa, despite its distant evolutionary placement within the Erysipelotrichia. The discovery of nutrient-responsive germination mechanisms, heat-activation dependence, and a Bacilli-style SpoVA-driven Ca–DPA release pathway highlights the conservation of canonical germination pathways in gut commensal.

## FUNDING

This work was supported by ANID/BASAL/FB210008 to R.Q., ANID/PhD Scholarship 21241350 to C.R-V. Awards from ANID – Millennium Science Initiative Program — NCN17_093, Texas A&M University Start Up funds, and 5R01AI177842 from the National Institute of Allergy and Infectious Diseases, all to D.P-S.

### DATA AVAILABILITY

This Whole Genome Shotgun project has been deposited at DDBJ/ENA/GenBank under the accession JAGBKO000000000. The version described in this paper is version JAGBKO010000000.

